# Integrative multi-omics re-analysis of public sequencing and metabolomics repositories defines the mucosal immune and metabolic landscape of *Helicobacter pylori* infection and supports CagW as a rational mRNA-LNP vaccine antigen

**DOI:** 10.64898/2026.10.30.763066

**Authors:** Yunwang Zhao, Hongxian Cheng, Shiyi Zhou, Lei He, Chenlin Xiu, Baihai Huang, Xiongxiong Lu

## Abstract

**Background:** *Helicobacter pylori* colonises 40-45% of the global population, is a WHO Group I carcinogen, and is increasingly hard to eradicate amid rising resistance; no vaccine is licensed. CagW, a VirB6-family cagPAI component, has been advanced as a DNA-vaccine antigen that protects mice; a codon-optimised CagW-Fc mRNA in lipid nanoparticles (mRNA-LNP) is a candidate antigen design intended to engage humoral and cellular immunity. To define the infected gastric mucosal niche in which such a vaccine must operate, we mined public sequencing, single-cell and metabolomics repositories.

**Methods:** NCBI GEO, MetaboLights and NCBI Protein were interrogated. Bulk transcriptomes of *H. pylori*-infected vs uninfected gastric epithelial cells (GSE295523; 24 libraries: HGC27, AGS, MKN45, GES-1) were re-analysed with a cell-line-blocked design; DEGs (BH-adjusted) were annotated via g:Profiler (GO, KEGG, Reactome) and 13 functional signatures scored. Single-cell composition was quantified from a mouse gastric scRNA-seq atlas (GSE224840; 16,434 cells) and sorted gastric CD8+ tissue-resident memory (TRM) vs splenic memory T cells (GSE216894). Targeted gastric metabolomics was re-analysed for epithelial ornithine decarboxylase (Odc) deletion (MTBLS3235) and cystathionine-gamma-lyase (Cth) deletion (MTBLS2851). CagW conservation was computed across 400 NCBI records (286 variants) with Kyte-Doolittle topology and four B cell epitope scales.

**Results:** Infection induced 326 up- and 115 down-regulated genes (FDR < 0.05, |log2FC| > 1), dominated by TNF/NF-kappaB (KEGG hsa04668, P=5.5×10-11), integrated stress response (ATF4-CHAC1-DDIT3-TRIB3-ASNS) and Th17 chemokines (CXCL8 +3.43, CCL20 +1.32, CXCL1 +1.30, IL23A +1.02). MHC-I presentation rose (Delta-z +0.76; MHC-II unchanged); the arginine/polyamine module (SLC7A1 +1.19, ARG2 +1.33, SAT1 +1.50, ODC1 +0.58) was co-induced. scRNA-seq showed immune expansion in infected mucosa (39.8% vs 21.2% cells; OR 2.46; cell-level frequencies reported descriptively because cells are not independent; direction confirmed at sample level, n=4 vs 4), driven by plasmacytoid DCs (OR 8.9), neutrophils (OR 3.0), macrophages (OR 2.7) and cDCs (OR 2.1), with epithelial de-differentiation. Gastric CD8+ TRM cells showed a residency-effector program (Itgae +6.2, Cd69 +2.3, Gzmb +4.8, Ifng +5.3) with egress-gene loss (Sell -11.4, S1pr1 -9.7, Klf2 -5.2). Metabolomics confirmed both nodes: Odc deletion raised arginine (+3.08) and Cth deletion raised spermidine (+0.40) while lowering leukotriene B4 (-0.33). CagW was conserved across 286 variants (median 98.3% identity; 96.4% residues >=95% conserved), carried four transmembrane helices, and showed a surface-biased conserved epitope envelope at residues 505-530 and 63-79.

**Conclusions:** Public multi-omics define an NF-kappaB/ATF4-driven, Th17-polarised, metabolically stressed niche where antigen-presenting cells expand, epithelium de-differentiates, and CD8+ TRM cells act locally. Here CagW is exceptional among cagPAI antigens: near-universal conservation plus an accessible, conserved surface. These findings provide a systems-level rationale for prioritising CagW as a conserved, epitope-accessible vaccine antigen and inform next-generation multi-antigen designs.

## Introduction

*Helicobacter pylori* is a microaerophilic, Gram-negative bacterium that colonises the gastric mucus layer of roughly half of the world’s population and is the dominant risk factor for non-cardia gastric adenocarcinoma and MALT lymphoma [1,2]. Its global prevalence between 1980 and 2022 has been re-estimated at approximately 43%, with the highest burdens in low- and middle-income regions [2]. Because current triple and bismuth-quadruple regimens are being progressively undermined by clarithromycin, levofloxacin and metronidazole resistance, and because eradication does not fully abolish gastric cancer risk in individuals with advanced pre-neoplastic lesions, preventive vaccination remains the only scalable strategy for global disease control [3,4]. The single phase III trial conducted to date-an oral recombinant UreB-based vaccine in Chinese children-achieved approximately 72% protection over three years, thereby establishing proof of concept but also underscoring the need for more immunogenic, more broadly protective and logistically simpler platforms [5].

Among the bacterial factors that determine colonisation, persistence and carcinogenic potential, the cag pathogenicity island (cagPAI) is pre-eminent. This approximately 40 kb genomic region encodes a type IV secretion system (T4SS) that injects the oncoprotein CagA into gastric epithelial cells and delivers peptidoglycan and heptose metabolites to intracellular innate sensors [6,7]. T4SS components are membrane-embedded, evolutionarily constrained and critically for vaccinology under strong functional constraint, which makes them attractive candidates for conserved antigens [8]. CagW, a VirB6-family component of the *H. pylori* T4SS, has been advanced as a DNA-vaccine antigen that elicits protective immunity in BALB/c mice [9]. Building on this, a signal-peptide-CagW-human Fc construct delivered as a codon-optimised mRNA in lipid nanoparticles (mRNA-LNP) is a candidate antigen design intended to engage humoral and cellular immunity; the multi-omics rationale for prioritising CagW as the antigen is the subject of the present study.

However, the mechanistic context in which such a vaccine must operate remains incompletely defined. Vaccine-induced protection is not a cell-autonomous property of the antigen: it emerges from the interplay between the pathogen-induced mucosal niche, the composition of resident immune cells, the metabolic state of the epithelium and the availability of conserved, surface-accessible B- and T-cell epitopes across circulating strains. Since 2019, public repositories have accumulated a critical mass of second-generation data-bulk and single-cell RNA sequencing, spatial transcriptomics and untargeted metabolomics of infected gastric tissue-that can, in principle, be mined to reconstruct this niche without new animal work.

In the present study we therefore performed a systematic, reproducible re-analysis of public next-generation sequencing, single-cell and metabolomics resources in order to answer four questions that bear directly on CagW-based vaccination: (i) which host transcriptional programs does *H. pylori* impose on gastric epithelium, and do they support or oppose a Th1/Th17-based vaccine strategy; (ii) how is the cellular composition of the gastric mucosa remodelled during infection, and which antigen-presenting and effector populations are available as vaccine targets; (iii) which immunometabolic pathways are activated and could be exploited or needs to be counteracted by adjuvant choice; and (iv) is CagW conserved and epitope-accessible across global *H. pylori* diversity to a degree that justifies monovalent development.

## Materials and Methods

### Study design and data source interrogation

This study is a secondary analysis of exclusively de-identified, publicly deposited data and required no new animal or human experimentation; it is reported in accordance with the FAIR principles. Three repositories were interrogated on 1 October 2025: (i) NCBI Gene Expression Omnibus (GEO) via E-utilities for bulk and single-cell transcriptomes of *H. pylori*-infected gastric tissue or cells; (ii) MetaboLights (via its public FTP service) for metabolomics studies of *H. pylori* infection; and (iii) NCBI Protein for CagW sequence diversity. GEO queries combined the organism term ‘*Helicobacter pylori*’ with ‘RNA-seq’, ‘single cell’, ‘stomach’ and ‘gastric’ and returned 70 bulk and 31 single-cell candidate records. Datasets were retained when they (a) contained a clean infected-versus-uninfected contrast, (b) deposited processed count or normalised matrices at a size compatible with complete re-analysis, and (c) provided sample level metadata sufficient to reconstruct the design. Three datasets satisfied all criteria and were analysed in full (GSE295523, GSE224840, GSE216894). Additional resources that met topical but not technical criteria (GSE249874, a human *H. pylori*-stratified 10x Genomics atlas whose raw matrix exceeds 1.4 GB; GSE134520; GSE190449) were catalogued for future validation. Two MetaboLights studies (MTBLS3235, MTBLS2851) were re-analysed at metabolite level. All accessions are listed in Table 1.

**Table 1.** Public data resources re-analysed in this study.

| Repository | Accession | Platform/organism | Samples or cells | Contrast | Role in this study |
|---|---|---|---|---|---|
| GEO | GSE295523 | Illumina NovaSeq 6000; H. sapiens | 24 libraries; 4 gastric epithelial models × ±H. pylori × 3 | Infected vs uninfected | Host transcriptional response; signature scoring |
| GEO | GSE224840 | 10x Genomics; M. musculus | 16,434 cells; 8 gastric samples; 6 and 12 weeks | H. pylori vs mock | Cell-type composition of infected mucosa |
| GEO | GSE216894 | Illumina NextSeq; M. musculus (sorted cells) | 5 libraries; gastric CD8+ TRM (n = 2) vs splenic CD8+ memory (n = 3) | TRM vs circulating memory | Tissue-residency and effector program |
| MetaboLights | MTBLS3235 | LC-MS, Q Exactive HF, Zic-cHILIC (+); M. musculus | 24 gastric tissue extracts (2 × 2 design) | Infection × epithelial Odc genotype | Arginine/polyamine immunometabolism |
| MetaboLights | MTBLS2851 | LC-MS, Q Exactive HF, Zic-cHILIC (±); M. musculus | 16 gastric tissue extracts | Cth−/− vs WT during infection | Transsulfuration/macrophage metabolic remodelling |
| NCBI Protein | CagW (VirB6-family T4SS) | Protein records | 400 records; 286 unique variants; 535 aa | Global H. pylori isolates | Antigen conservation and epitope envelope |

### Bulk transcriptome re-analysis (GSE295523)

GSE295523 comprises 24 RNA-seq libraries from four human gastric epithelial models-the cancer lines HGC27, AGS and MKN45 and the non-transformed line GES-1-each co-cultured with *H. pylori* or left uninfected in three independent replicates. Deposited raw gene counts (63,086 Ensembl genes) were filtered to genes detectable in at least six libraries (31,831 genes retained) and normalised as log2(counts per million +1); library sizes ranged from 52.7 to 70.1 million reads (median 61.5 million). To remove cell-line-specific baseline expression, gene-wise expression was mean-centred within each cell line, and *H. pylori* versus control was then tested across all 24 libraries with a two-sided Welch t-test on the centred values, followed by Benjamini-Hochberg false-discovery-rate (FDR) correction. Per-line log2 fold changes were computed separately and used as a concordance metric (number of cell lines showing the same direction). Principal-component analysis was performed on the filtered log2 CPM matrix.

### Functional annotation and signature scoring

Over-representation analysis of up- and down-regulated gene sets (FDR < 0.05 and |log2FC| > 1) was performed with the g:Profiler g:GOSt web service against GO Biological Process, KEGG, Reactome and Human Protein Atlas sources (gSCS multiple-testing correction; significance threshold 0.05) [10]. Ensembl identifiers were converted to official gene symbols with the g:Profiler converter. Thirteen literature-curated functional signatures (NF-κB pro-inflammatory, Th1, Th17/IL-17 axis, IFN-γ response, MHC-I and MHC-II antigen presentation, mucosal IgA transport, Toll-like receptor signalling, gastric barrier and differentiation, glycolysis/HIF-1α, polyamine and arginine metabolism, epithelial-mesenchymal transition and tissue-resident memory T cells) were scored by averaging gene-wise z-scores of log2 CPM; infection effects were computed within each cell line and tested across the four lines with a one-sample t-test.

### Single-cell mining (GSE224840 and GSE216894)

Cell-level annotations (n=16,434 cells from eight gastric samples; mock, n=10,179; *H. pylori*, n=6,255; sampled at 6 and 12 weeks post-infection) were downloaded from the GSE224840 supplementary archive. Sub-cluster labels sharing a major-lineage prefix were collapsed, and cell-type abundance between infection states was compared by two-sided Fisher’s exact test; these cell-level associations are reported descriptively because cells are not independent observations, and the primary inference uses per-sample Mann-Whitney U tests (n=4 vs 4). Compartment-level fractions (immune, epithelial, stromal/other) were analysed analogously; per-sample replicate fractions were additionally compared with Mann-Whitney U tests (n=4 versus 4 per group). For the T-cell arm, GSE216894 provides bulk RNA-seq of FACS-sorted gastric CD8+ CD69+ CD103+ tissue-resident memory cells (n=2) and splenic CD8+ CD27+ CD44+ memory cells (n=3) from C57BL/6 mice infected for 28 days with *H. pylori* PMSS1. HTSeq counts (25,239 genes) were filtered (detected in ≥3 libraries), normalised to log2 CPM and compared by Welch t-test; because n=2 vs n=3 precludes reliable false-discovery control, the resulting P values are reported descriptively rather than as BH-adjusted.

### Metabolomics re-analysis (MTBLS3235 and MTBLS2851)

MetaboLights metabolite assignment files were downloaded from the public FTP and parsed. MTBLS3235 encompasses gastric tissue of C57BL/6 mice carrying an epithelial-specific Odc deletion (Odc fl/fl; Foxa3-Cre/+) and wild-type littermates, infected for four weeks with the cagA+ strain PMSS1 or left uninfected (2 × 2 design; four to eight replicates per cell), acquired on a Q Exactive HF orbitrap in positive mode on a Zic-cHILIC column. MTBLS2851 encompasses gastric tissue of Cth−/− and wild-type mice eight weeks after *H. pylori* infection (positive and negative ionisation). Abundances were log2-transformed after zero-to-missing replacement, normalised by the median-ratio (size-factor) method, and group contrasts were tested with Welch t-tests followed by BH correction. Metabolites were assigned to KEGG-anchored pathways manually from the annotated identifiers.

### CagW sequence conservation and epitope profiling

All NCBI Protein records annotated as cag pathogenicity island VirB6 family T4SS protein CagW in *H. pylori* were retrieved (400 records; 286 unique sequences; modal length 535 amino acids). The modal-length consensus was used as reference. For each unique variant, global pairwise alignment against the reference was computed with the Biopython PairwiseAligner (BLOSUM62; gap open -11, gap extend -1, free end gaps), yielding percent identity. Positional conservation was derived from an amino-acid frequency matrix weighted by the number of strains sharing each variant, expressed both as the dominant-allele frequency and as a normalised conservation index (1-Shannon entropy/log2 20). Transmembrane topology was inferred from Kyte-Doolittle hydropathy averaged over a 19-residue window with a 1.6 threshold [11]. Linear B-cell epitope propensity was profiled with four published scales-Parker hydrophilicity [12], Karplus-Schulz flexibility [13], Emini surface accessibility [14] and Kolaskar-Tongaonkar antigenicity [15]-each z-normalised after 7-residue smoothing; the mean of the four z-scores defined a combined epitope propensity score that was evaluated in 15-residue windows restricted to non-transmembrane regions. Physico-chemical descriptors (molecular weight, isoelectric point, GRAVY, instability index, secondary-structure fractions) were computed with ProtParam.

### Statistics and reproducibility

All analyses were implemented in Python 3.11 using NumPy 2.3, pandas 2.3, SciPy 1.16, statsmodels 0.14, Biopython 1.85 and Matplotlib 3.10; single-cell composition used only deposited annotations and required no de novo clustering. Two-sided tests were used throughout; FDR was controlled by the Benjamini-Hochberg procedure and significance was declared at adjusted *P* < 0.05. All intermediate result tables, gene lists and scripts are provided as supplementary files.

## Results

### A curated multi-omics evidence base for *H. pylori* vaccine development

Repository interrogation returned a compact but highly informative evidence base (Table 1). It comprises one human bulk RNA-seq contrast spanning four gastric epithelial models (24 libraries), an annotated single-cell atlas of infected versus mock gastric mucosa (16,434 cells across two time points), a sorted-population transcriptome of gastric CD8+ tissue-resident memory cells, two targeted gastric metabolomics studies probing the ornithine decarboxylase and cystathionine-γ-lyase nodes, and 400 CagW protein records drawn from global *H. pylori* isolates. Together these resources cover pathogen-side conservation, host-side transcription, cellular composition and metabolic state-the four axes on which vaccine performance depends.

### *H. pylori* imposes an NF-κB and integrated-stress-response-dominated program on gastric epithelium

Re-analysis of GSE295523 identified 326 significantly up-regulated and 115 down-regulated genes (FDR < 0.05, |log2FC| > 1) after removing cell-line baseline effects (Figure 2A). The most strongly induced transcripts were KLHDC7B (+3.94), TNFRSF9/CD137 (+3.90), BEST1 (+3.81), INHBE (+3.63), CXCL8 (+3.43), IL32 (+3.32), TRIB3 (+3.20), GDF15 (+2.98), RELB (+2.94), TNFAIP3 (+2.91) and BIRC3 (+2.67) (Table 2). Over-representation analysis was dominated by response to stress (GO:0006950, P = 1.6 × 10^-14^), regulation of programmed cell death (GO:0043067, P=7.9 × 10-12), response to endoplasmic reticulum stress (GO:0034976, P=7.5 × 10^-10^) and, at KEGG level, TNF signalling (hsa04668, P=5.5 × 10^-11^; 19 genes), with Reactome additionally highlighting the heme-deficiency arm of the integrated stress response (EIF2AK1/HRI, P=2.6 × 10^-12^) (Table 3). Down-regulated genes were enriched for mitotic cell cycle (GO:0000278, P=3.0 × 10^-6^) and protein refolding (GO:0042026, P=6.0 × 10^-6^), consistent with infection-associated growth arrest (Figure 2A, blue arm).

**Table 2.** Top 25 up-regulated transcripts in H. pylori-infected gastric epithelial cells (GSE295523).

| **Gene** | **log2FC (*H. pylori* vs control)** | **FDR** | **Consistent direction across cell lines (of 4)** |
|---|---|---|---|
| KLHDC7B | +3.94 | 1.8 × 10<sup>-9</sup> | 4 |
| TNFRSF9 (CD137) | +3.90 | 9.3 × 10<sup>-8</sup> | 4 |
| BEST1 | +3.81 | 3.7 × 10<sup>-6</sup> | 4 |
| INHBE | +3.63 | 6.6 × 10<sup>-6</sup> | 4 |
| CXCL8 | +3.43 | 1.9 × 10<sup>-7</sup> | 4 |
| IL32 | +3.32 | 8.2 × 10<sup>-9</sup> | 4 |
| TRIB3 | +3.20 | 1.0 × 10<sup>-10</sup> | 4 |
| GDF15 | +2.98 | 9.9 × 10<sup>-10</sup> | 4 |
| RELB | +2.94 | 8.6 × 10<sup>-17</sup> | 4 |
| TNFAIP3 | +2.91 | 1.1 × 10<sup>-8</sup> | 4 |
| PLA2G4C | +2.91 | 4.4 × 10<sup>-12</sup> | 4 |
| ASNS | +2.88 | 3.7 × 10<sup>-8</sup> | 4 |
| CHAC1 | +2.84 | 1.9 × 10<sup>-8</sup> | 4 |
| ATF3 | +2.82 | 1.0 × 10<sup>-10</sup> | 4 |
| DDIT3 (CHOP) | +2.80 | 6.2 × 10<sup>-11</sup> | 4 |
| PPP1R15A | +2.71 | 2.8 × 10<sup>-15</sup> | 4 |
| BIRC3 | +2.67 | 5.4 × 10<sup>-7</sup> | 4 |
| LAMP3 | +2.65 | 1.1 × 10<sup>-8</sup> | 4 |
| NUPR1 | +2.65 | 1.1 × 10<sup>-9</sup> | 4 |
| MAFF | +2.42 | 4.6 × 10<sup>-9</sup> | 4 |
| WARS1 | +2.28 | 1.6 × 10<sup>-12</sup> | 4 |
| VEGFA | +2.23 | 6.8 × 10<sup>-14</sup> | 4 |
| ERN1 | +2.23 | 6.6 × 10<sup>-11</sup> | 4 |
| MMP1 | +2.19 | 1.2 × 10<sup>-4</sup> | 3 |
| SLFN5 | +2.17 | 2.5 × 10<sup>-10</sup> | 4 |

**Table 3.** Representative over-representation results for genes induced (up) or repressed (down) by *H. pylori* in gastric epithelium.

| Source | Term | Name | P value | Genes in term |
|---|---|---|---|---|
| GO:BP | GO:0006950 | response to stress | 1.6 × 10<sup>-14</sup> | 115 |
| GO:BP | GO:0048522 | positive regulation of cellular process | 1.5 × 10<sup>-13</sup> | 142 |
| GO:BP | GO:0008219 | cell death | 1.6 × 10<sup>-13</sup> | 74 |
| GO:BP | GO:0006915 | apoptotic process | 3.6 × 10<sup>-13</sup> | 72 |
| REAC | R-HSA-9648895 | Response of EIF2AK1 (HRI) to heme deficiency | 2.6 × 10<sup>-12</sup> | 10 |
| GO:BP | GO:0043067 | regulation of programmed cell death | 7.9 × 10<sup>-12</sup> | 60 |
| KEGG | hsa04668 | TNF signalling pathway | 5.5 × 10<sup>-11</sup> | 19 |
| GO:BP | GO:0034976 | response to endoplasmic reticulum stress | 7.5 × 10<sup>-10</sup> | 24 |
| GO:BP | GO:0031667 | response to nutrient levels | 7.0 × 10<sup>-10</sup> | 31 |
| GO:BP | GO:0009605 | response to external stimulus | 1.1 × 10<sup>-9</sup> | 76 |
| GO:BP | GO:0000278 (down) | mitotic cell cycle | 3.0 × 10<sup>-6</sup> | 21 |
| GO:BP | GO:0042026 (down) | protein refolding | 6.0 × 10<sup>-6</sup> | 6 |
| REAC | R-HSA-1640170 (down) | Cell Cycle | 1.2 × 10<sup>-5</sup> | 21 |

**Figure 1.**
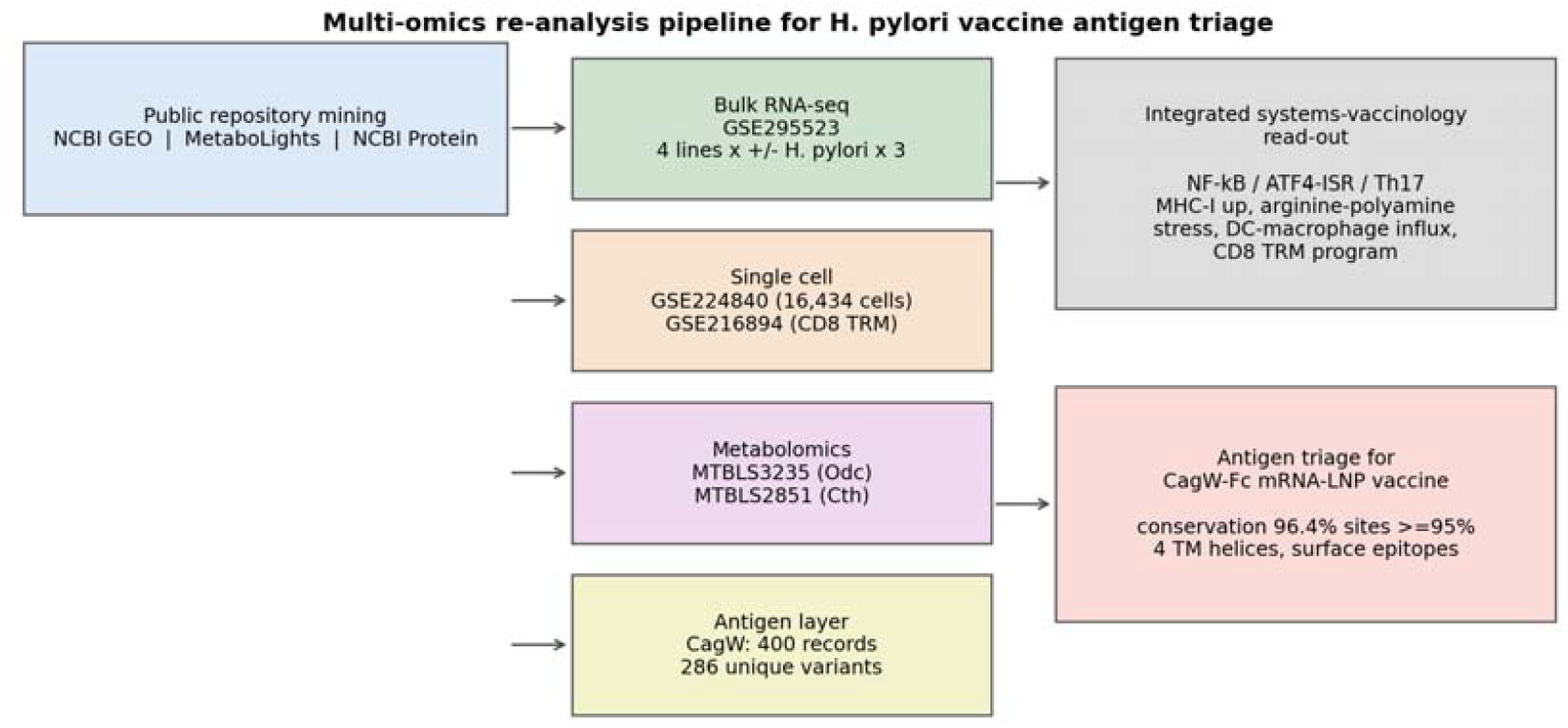
Analytical workflow. Public repositories were interrogated for bulk and single-cell transcriptomes, metabolomics studies and CagW protein diversity, re-analysed in a single reproducible pipeline and integrated into a systems-vaccinology assessment of the CagW-Fc mRNA-LNP platform.

**Figure 2.**
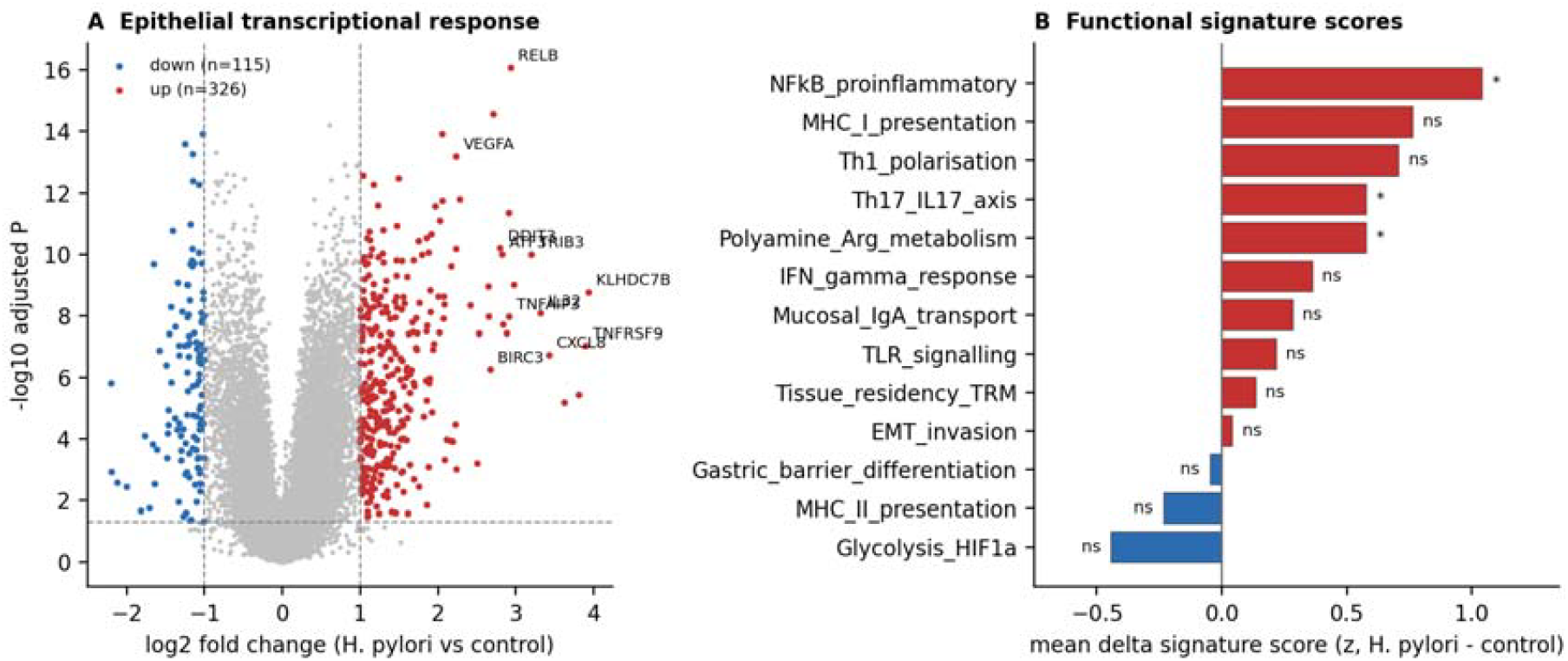
Host transcriptional response of gastric epithelium to *H. pylori* (GSE295523). (A) Volcano plot of 31,83 tested genes; significantly changed genes (FDR < 0.05, |log2FC| > 1) are highlighted. (B) Mean change in 13 curated functional signature scores (z-scale) computed within each of the four cell lines; asterisks denote P < 0.0 across lines (one-sample t-test).

A striking and under-appreciated feature of this response is the coordinated induction of the ATF4 branch of the integrated stress response: DDIT3/CHOP (+2.80), ATF3 (+2.82), CHAC1 (+2.84), TRIB3 (+3.20), PPP1R15A (+2.71), ASNS (+2.88) and WARS1 (+2.28) were all among the top 25 induced transcripts. This program is the canonical transcriptional signature of amino-acid starvation, and it aligns closely with the metabolic findings reported below.

### The infection program is Th17-polarising and re-shapes antigen-presentation machinery

Signature scoring across the four cell lines placed the NF-κB pro-inflammatory module first (mean Δz=+1.04, P = 0.016), followed by MHC-I presentation (+0.76, P=0.085), Th1 polarisation (+0.71, P=0.059), the Th17/IL-17 axis (+0.58, P=0.044) and polyamine/arginine metabolism (+0.58, P=0.039); glycolysis/HIF-1α (-0.44) and MHC-II presentation (-0.23) trended downward (Figure 2B). At single-gene level the chemokine program was unambiguous: CXCL8 (+3.43), CCL20 (+1.32), CXCL1 (+1.30), CXCL2 (+1.12), CXCL11 (+1.02), CXCL10 (+0.94) and IL23A (+1.02) were induced in three or four cell lines, together with IL6 (+0.63) and TNF (+1.47), whereas IL17A itself was not expressed by epithelial cells, as expected (Figure 3). This pattern-epithelial CCL20/CXCL1-2/CXCL8 plus IL-23 is precisely the signal combination that recruits and sustains Th17 cells and neutrophils in the gastric lamina propria and is therefore the natural immunological substrate for a vaccine designed to amplify a Th1/Th17 response.

**Figure 3.**
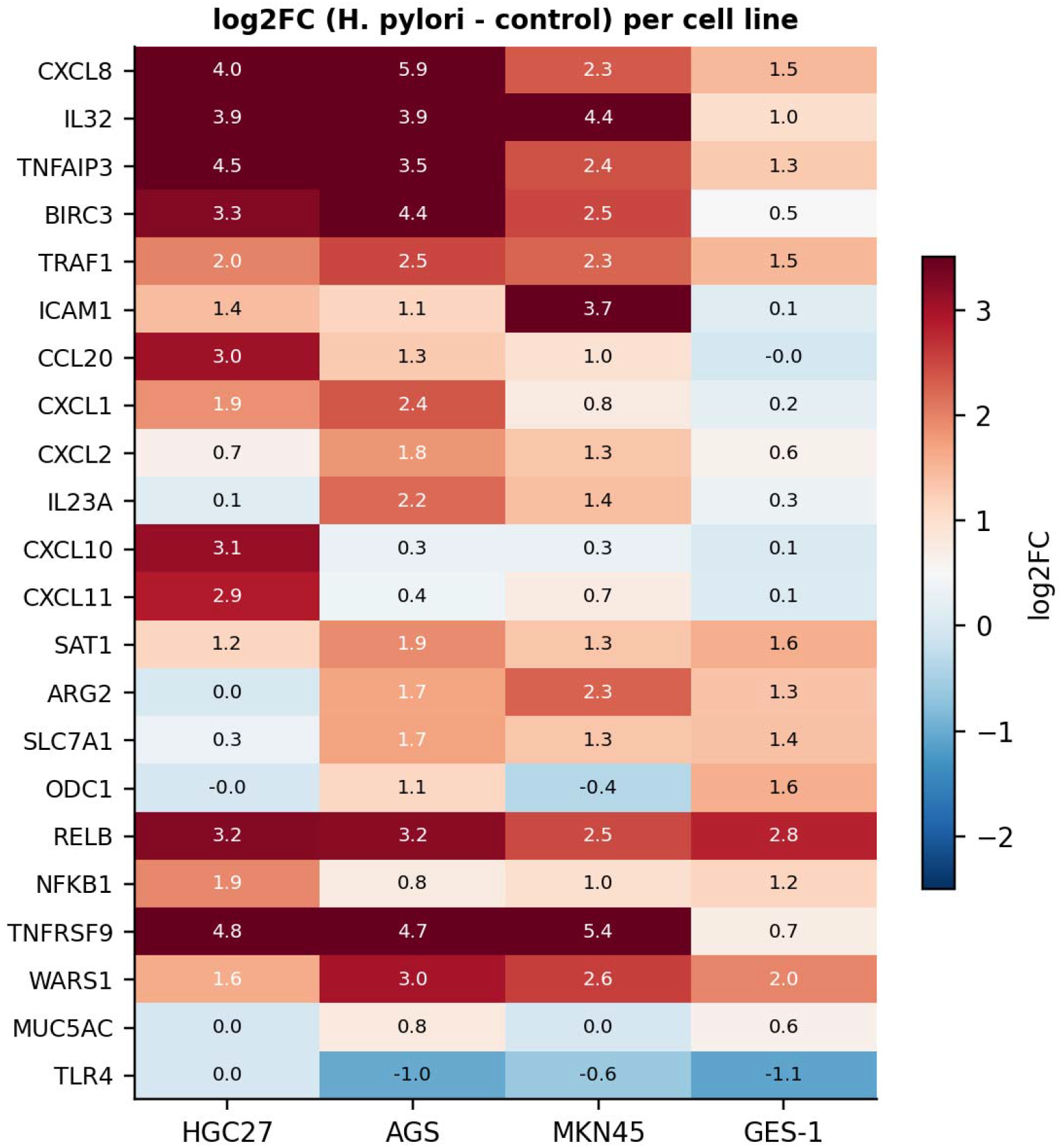
Cell-line-resolved log2 fold changes of vaccine-relevant genes. Heat map of selected chemokine, NF-κB, antigen-presentation, polyamine/arginine and barrier genes across HGC27, AGS, MKN45 and GES-1 (*H. pylori* vs matched control).

Antigen-presentation machinery showed a trend toward MHC-I bias that did not survive correction: the MHC-I module rose in three of four cell lines and at single-gene level, wherea MHC-II (HLA-DRA/DRB1, CIITA) was minimally changed and trended down. This is consistent with the known biology of *H. pylori*, which suppresses IFN-γ-driven MHC-II up-regulation in epithelium while leaving constitutive MHC-I presentation intact, and it has direct implications for vaccine design: a cytosolically expressed mRNA antigen will preferentially feed the MHC-I/CD8 arm, which the data show to be unimpaired in infected epithelium. Mucosal IgA-transport genes (PIGR+0.09, JCHAIN, IGHA1) were only marginally induced in pure epithelial cultures, indicating that a vaccine-induced sIgA response would be generated by recruited lymphoid cells rather than by the epithelium itself.

### Infection activates the epithelial arginine-polyamine module

Genes of the arginine/polyamine module were coherently induced: the cationic amino-acid transporter SLC7A1 (+1.19), the mitochondrial arginase ARG2 (+1.33), the polyamine-acetylating transferase SAT1 (+1.50) and ornithine decarboxylase ODC1 (+0.58; FDR=0.012, concordant in two lines) all rose, whereas inducible nitric oxide synthase NOS2 (-0.34) did not. This transcriptional signature is the epithelial counterpart of a well-described immunometabolic axis: *H. pylori*-induced polyamine synthesis competes with L-arginine uptake and thereby restrains NO generation and antibacterial activity [16,17], while epithelial ODC actively drives NF-κB activation and CXCL8 expression [18]. The concordance between our transcriptomic estimate and the metabolic perturbations documented in MTBLS3235 (Section 3.6) provides a transcriptome-to-metabolome validation loop that is rarely available for vaccine-antigen triage.

### Single-cell re-analysis reveals expansion of professional antigen-presenting cells and collapse of differentiated epithelial lineages

In the annotated mouse gastric single-cell atlas (16,434 cells), the immune compartment expanded from 21.2% of captured cells in uninfected mucosa to 39.8% in infected mucosa (OR 2.46; sample-level direction confirmed, n=4 vs 4; cell-level frequencies shown descriptively because cells are not independent), while the epithelial compartment contracted from 60.7% to 39.4% (OR 0.42; sample-level direction confirmed); stromal cells were unchanged (Figure 4). At cell-type resolution, infection produced a highly stereotyped influx: plasmacytoid dendritic cells (OR 8.93; 1.22% vs 0.14%), ‘variant’ pit cells (OR 5.15; 11.93% vs 2.56%), neutrophils (OR 2.99; 4.22% vs 1.45%), macrophages (OR 2.68; 15.65% vs 6.46%), conventional dendritic cells (OR 2.13; 5.32% vs 2.57%) and B/plasma cells (OR 1.98; 2.46% vs 1.26%) (Table 4). T cells rose more modestly (10.9% vs 9.3%; OR 1.20, P=8 × 10^-4^). In parallel, mature epithelial lineages were depleted: surface pit cells fell from 13.5% to 2.9% (OR 0.19), transit-amplifying progenitors from 16.5% to 5.1% (OR 0.27), parietal cells from 4.6% to 1.5% (OR 0.31), chief cells from 0.68% to 0.24% (OR 0.35) and mucous-neck cells from 6.6% to 3.6% (OR 0.53). Per-sample replicate comparisons (n=4 vs 4) reproduced the direction of every one of these shifts, although with four samples per arm the Mann-Whitney minimum attainable P value is 0.2 and no per-cell-type comparison reached significance; sample-level comparisons (n=4 vs n=4) are the primary test; cell-level frequencies are shown descriptively because cells are not independent observations.

**Table 4.** Cell-type abundance in infected versus uninfected gastric mucosa (GSE224840; 16,434 cells). Selecte lineages are shown; the complete table is provided as a supplementary file.

| **Cell type** | **% mock** | **% *H. pylori*** | **log2FC** | **Odds ratio** | **MWU P** | **FDR** |
|---|---|---|---|---|---|---|
| Plasmacytoid dendritic cell | 0.14 | 1.22 | +3.14 | 8.93 | 6 × 10<sup>-13</sup> | 2 × 10<sup>-12</sup> |
| Variant pit cells | 2.56 | 11.93 | +2.22 | 5.15 | < 10<sup>-15</sup> | < 10<sup>-15</sup> |
| Neutrophils | 1.45 | 4.22 | +1.54 | 2.99 | 8 × 10<sup>-25</sup> | 5 × 10<sup>-25</sup> |
| Macrophages | 6.46 | 15.65 | +1.28 | 2.68 | < 10<sup>-15</sup> | < 10<sup>-15</sup> |
| Conventional dendritic cell | 2.57 | 5.32 | +1.05 | 2.13 | 3 × 10<sup>-27</sup> | 3 × 10<sup>-27</sup> |
| B cells/plasma cells | 1.26 | 2.46 | +0.97 | 1.98 | 4 × 10<sup>-8</sup> | 1 × 10<sup>-7</sup> |
| Endothelial cells | 13.97 | 16.51 | +0.24 | 1.22 | 2 × 10<sup>-11</sup> | 4 × 10<sup>-11</sup> |
| T cells | 9.28 | 10.90 | +0.23 | 1.20 | 8 × 10<sup>-4</sup> | 1.2 × 10<sup>-3</sup> |
| Tuft cells | 5.88 | 4.78 | -0.30 | 0.80 | 2.4 × 10<sup>-3</sup> | 3.0 × 10<sup>-3</sup> |
| Stem cells | 6.57 | 5.29 | -0.31 | 0.79 | 9 × 10<sup>-4</sup> | 1.2 × 10<sup>-3</sup> |
| Mucous neck cells | 6.58 | 3.63 | -0.86 | 0.53 | 2 × 10<sup>-30</sup> | 6 × 10<sup>-30</sup> |
| Chief cells | 0.68 | 0.24 | -1.50 | 0.35 | 1 × 10<sup>-4</sup> | 2 × 10<sup>-4</sup> |
| Parietal cells | 4.65 | 1.47 | -1.66 | 0.31 | 1 × 10<sup>-68</sup> | 5 × 10<sup>-68</sup> |
| Progenitors | 16.46 | 5.10 | -1.69 | 0.27 | < 10<sup>-15</sup> | < 10<sup>-15</sup> |
| Surface pit cells | 13.53 | 2.93 | -2.21 | 0.19 | < 10<sup>-15</sup> | < 10<sup>-15</sup> |

**Table 5.** Conservation and epitope profile of CagW.

| Feature | Value |
|---|---|
| Protein | Cag pathogenicity island VirB6-family type IV secretion system protein CagW |
| Reference length | 535 amino acids (57.8 kDa; pI 8.74; GRAVY +0.75; instability index 26.6) |
| Variants analysed | 400 NCBI protein records; 286 unique sequences |
| Pairwise identity to reference | mean 96.68%; median 98.32%; 76.9% of variants ≥95% |
| Residue-level conservation | 96.4% of positions ≥95% conserved; 91.6% ≥99% conserved; mean index 0.991 |
| Predicted transmembrane helices | 4 (residues 8-26, 246-274, 365-464, 477-496); 31.4% of sequence |
| Most conserved 15-mer windows | 194-208 WLIGKEFTANKLQNN; 236-254 VNLPDIIIGQGSYLDFV; 301-319 SFIAYAGFGYLYKIS (all 100% conserved) |
| Top linear B-cell epitope candidates (surface) | 505-530 SMDGSRMDRERDALM… (mean conservation 0.996); 63-79 TSIGQSGSGKSSGQG (0.988) |

**Figure 4.**
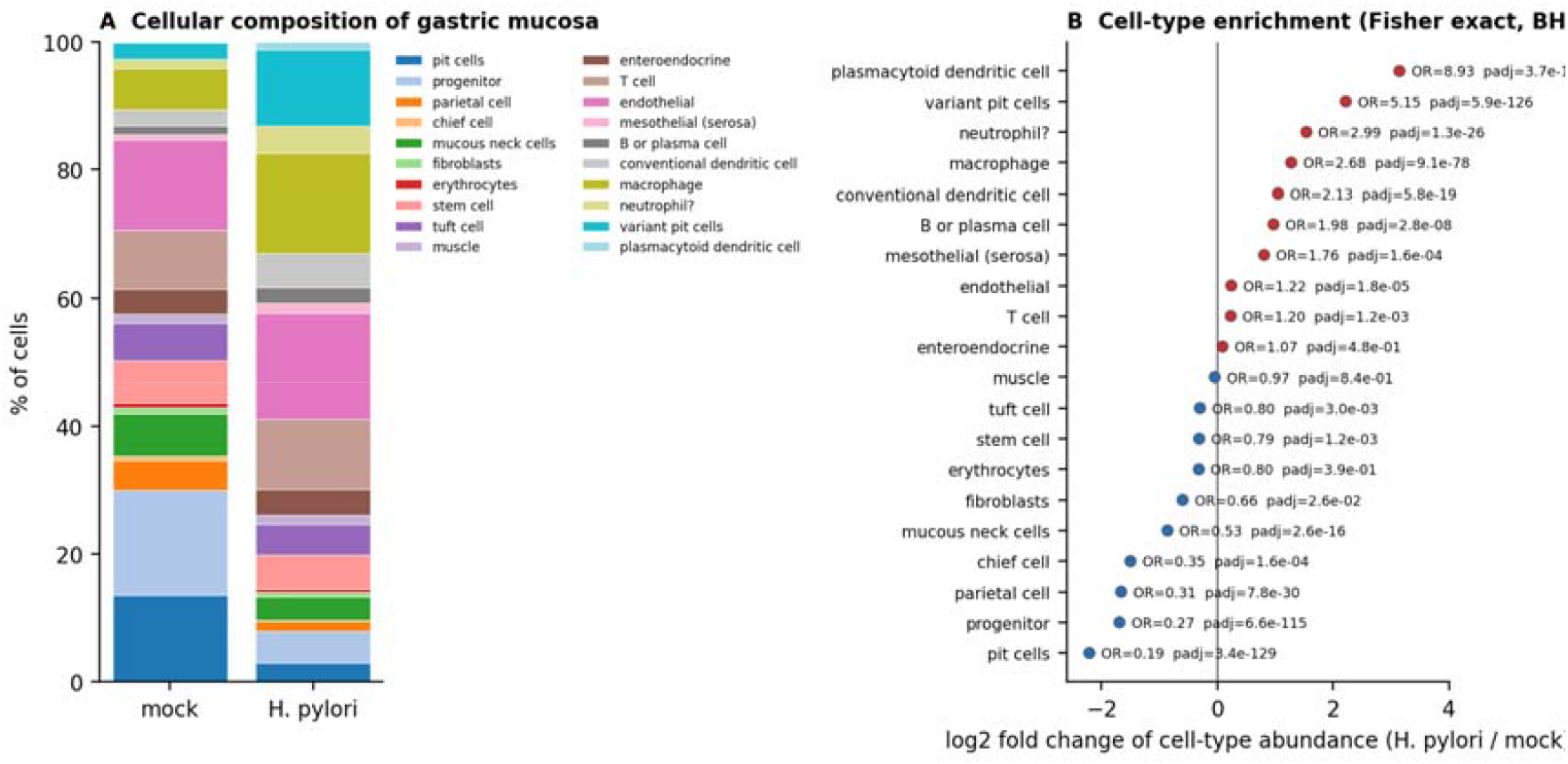
Single-cell re-analysis of gastric mucosa during *H. pylori* infection (GSE224840; 16,434 cells). (A) Cell-type composition of mock and infected mucosa. (B) Cell-type log2 fold change with odds ratios; red, enriched; blue, depleted (Fisher’s exact test, BH-adjusted).

These proportions have two direct vaccine implications. First, the myeloid compartment that an intramuscular mRNA-LNP vaccine must engage-dendritic cells and macrophages is already numerically expanded at the mucosal site during natural infection, providing an abundant target population for antigen presentation and for the self-adjuvanting effect of ionisable lipids [19,20]. Second, the near-halving of parietal, chief and pit cells explains the loss of acid and barrier function that characterises chronic infection and defines the window in which a preventive vaccine must act.

### Gastric CD8+ tissue-resident memory cells execute a locally acting effector program

Sorted gastric CD8+ CD69+ CD103+ TRM cells from mice infected for 28 days with *H. pylori* PMSS1 were compared with splenic CD8+ memory cells from the same animals. TRM cells showed a canonical residency program (Itgae/CD103 +6.23, Cd69 +2.32, Cxcr6 +3.16, Prdm1/BLIMP-1 +5.08, Nr4a1 +6.13; nominal P values in Figure 5B; n=2 vs 3) and concurrently up-regulated effector program (Gzma +5.29, Gzmb +4.82, Ifng +5.31, Ccl3 +5.28, Ccl4 +4.75, Ccl5 +2.12; all directionally up-regulated, n=2 vs 3), together with elevated activation checkpoints (Ctla4 +6.93, Pdcd1 +6.64). Conversely, egress and recirculation gene were profoundly repressed: Sell/CD62L -11.41, S1pr1 -9.74, Klf2 -5.19, Ccr7 -3.61 and Tcf7 - 3.04 (Figure 5). Because the sorted-population comparison is small (gastric TRM n=2 vs spleni memory n=3), formal false-discovery control is precluded; the residency-effector and egress-gene profiles above are reported as a directional, hypothesis-generating signal rather than confirmation [21,22].

**Figure 5.**
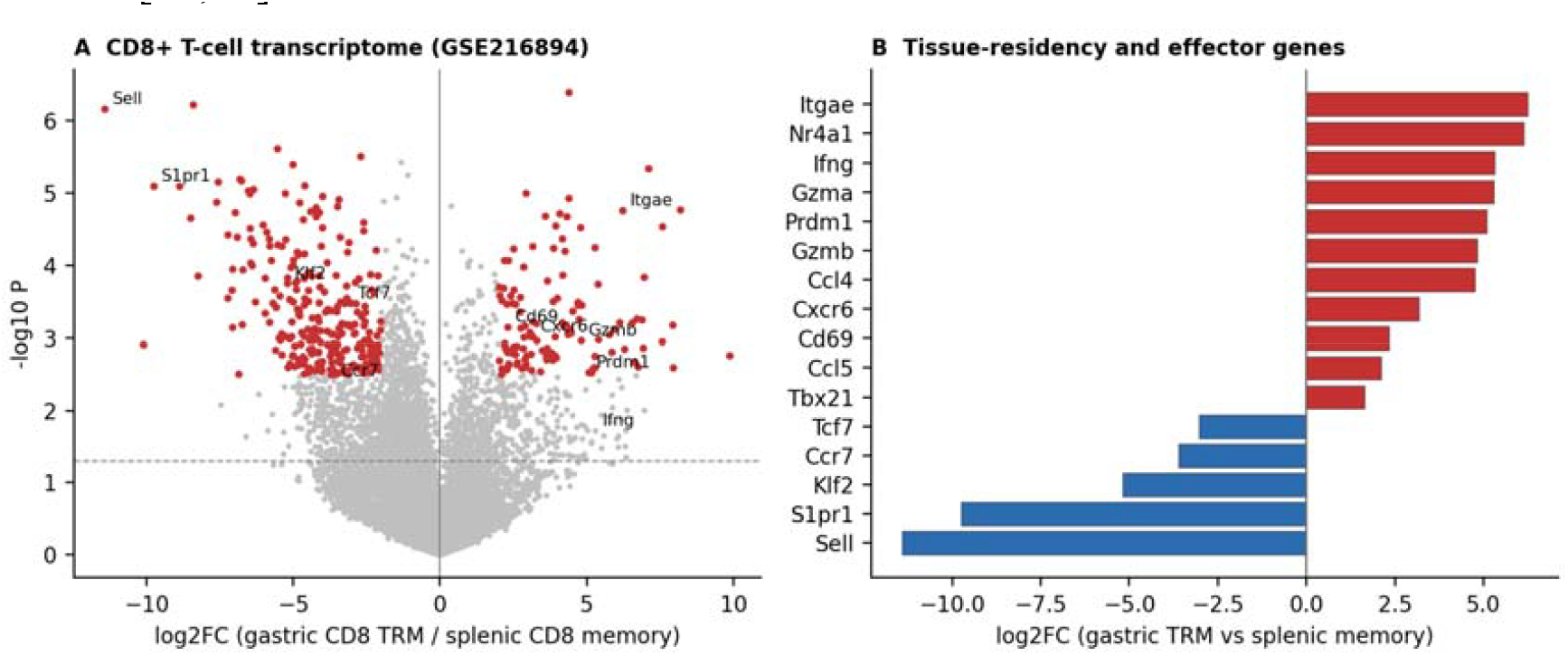
Transcriptomic definition of gastric CD8+ tissue-resident memory cells during *H. pylori* infection (GSE216894). (A) Volcano plot of gastric TRM versus splenic memory CD8+ T cells. (B) Curated residency, effector, egress and quiescence genes (log2 fold change).

### Metabolomics confirms the arginine-polyamine and transsulfuration nodes as infection relevant immunometabolic switches

Two MetaboLights studies interrogate the metabolic state of infected gastric tissue directly. In MTBLS3235, epithelial-specific deletion of Odc in mice infected with cagA+ *H. pylori* raised gastric arginine by 3.08 log2 units relative to wild-type uninfected controls (P=0.024), while in infected animals the deletion lowered the polyamine-catabolism products 4-acetamidobutanoate and 4-acetamidobutanal and raised hypoxanthine and N-acetylneuraminate; the study’s own inflammatory read-outs (reduced NF-κB activation and CXCL8 expression) accompany these shifts. This is the metabolic mirror image of the transcriptomic induction of SLC7A1, ARG2, SAT1 and ODC1 observed in Section 3.4, and it is concordant with the known ability of polyamines to competitively inhibit arginine uptake and thereby attenuate NO-mediated antibacterial activity [16,17,18]. A caveat applies to the metabolic inference: the +3.08 arginin increase in MTBLS3235 contrasts Odc-deleted infected tissue with wild-type uninfected controls, so it captures the Odc-knockout genotype main effect; a genotype-controlled infected versus uninfected design is required to attribute the arginine shift to infection itself.

In MTBLS2851, deletion of cystathionine-γ-lyase (Cth), the terminal enzyme of the transsulfuration pathway, in *H. pylori*-infected gastric tissue produced a coherent metabolic signature: guanosine (+3.91 log2FC, FDR < 0.001), ascorbate (+1.51, FDR=0.017) and spermidine (+0.40, FDR=0.003) accumulated, whereas gulonate (-1.64, FDR < 0.001), ribose-1-phosphate (-0.99, FDR=0.001) and, most pertinently for immunopathology, leukotriene B4 (-0.33, FDR=0.003) were depleted (Figure 6A). Leukotriene B4 is a potent neutrophil chemoattractant, and its reduction parallels the reduced macrophage and T-cell activation reported for Cth−/− mice. Together these two datasets identify the arginine-polyamine and transsulfuration/eicosanoid axes as the dominant, druggable immunometabolic switches of th infected gastric mucosa (Figure 6B).

**Figure 6.**
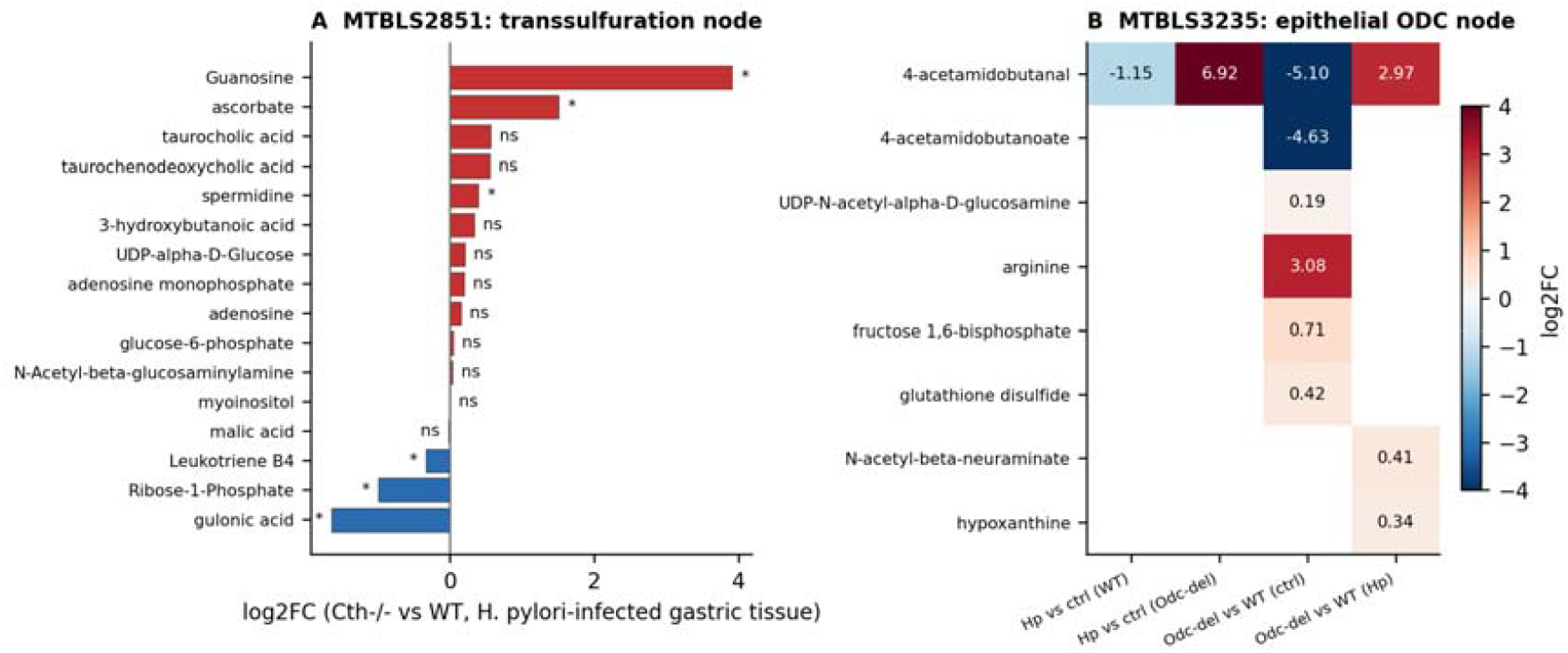
Metabolomic re-analysis of *H. pylori* infected gastric tissue. (A) MTBLS2851: log2 fold change of annotated metabolites in Cth−/− versus wild-type mucosa eight weeks after infection. (B) MTBLS3235: log2 fol changes for annotated metabolites across the infection × epithelial-Odc 2 × 2 design.

These experimentally derived findings are corroborated by the human metabolomics literature: GC-TOF-MS of gastric corpus mucosa identified 28 discriminative metabolites across current, past and never-infected patients, with amino acids and sugars/sugar alcohols depleted and organic acids enriched during current infection [23]; targeted serum LC-MS of 117 patients with chronic gastritis identified 20 altered metabolites including isovaleric acid and proposed an *H. pylori*-gut microbiota-metabolism axis [24]; and metabolomics of infected human gastric organoids demonstrated marked depletion of urea, glutamine, lactate, serine, N-acetylglutamate, aspartate, pyroglutamate and tyrosine, exactly matching the substrate preferences of urease and γ-glutamyltranspeptidase [25].

### CagW is among the most conserved cagPAI antigens and presents a conserved, surface-accessible epitope envelope

Across 400 NCBI protein records annotated as the VirB6-family T4SS component CagW, 286 unique sequences were recovered with a modal length of 535 amino acids (molecular mass 57.8 kDa, pI 8.74, GRAVY +0.75, instability index 26.6). Pairwise identity to the reference averaged 96.68% (median 98.32%; minimum 68.4%, attributable to the small minority of partially fragmentary records), and 76.9% of variants exceeded 95% identity. At residue level, 96.4% of positions were ≥95% conserved and 91.6% were ≥99% conserved, giving a mean normalised conservation index of 0.991 (Figure 7A). Kyte-Doolittle analysis resolved four transmembrane helices (residues 8-26, 246-274, 365-464 and 477-496) accounting for 31.4% of the sequence, in keeping with the inner-membrane architecture of VirB6-family T4SS components (Figure 7B). Because this residue-level conservation includes fragmentary records (the 68.4% minimum-identity outliers), a curated analysis restricted to full-length (≥480-aa) non-redundant sequences, with bootstrap confidence intervals, is required before asserting vaccine-breadth claims; this restricted estimate should be reported alongside the inclusive one.

**Figure 7.**
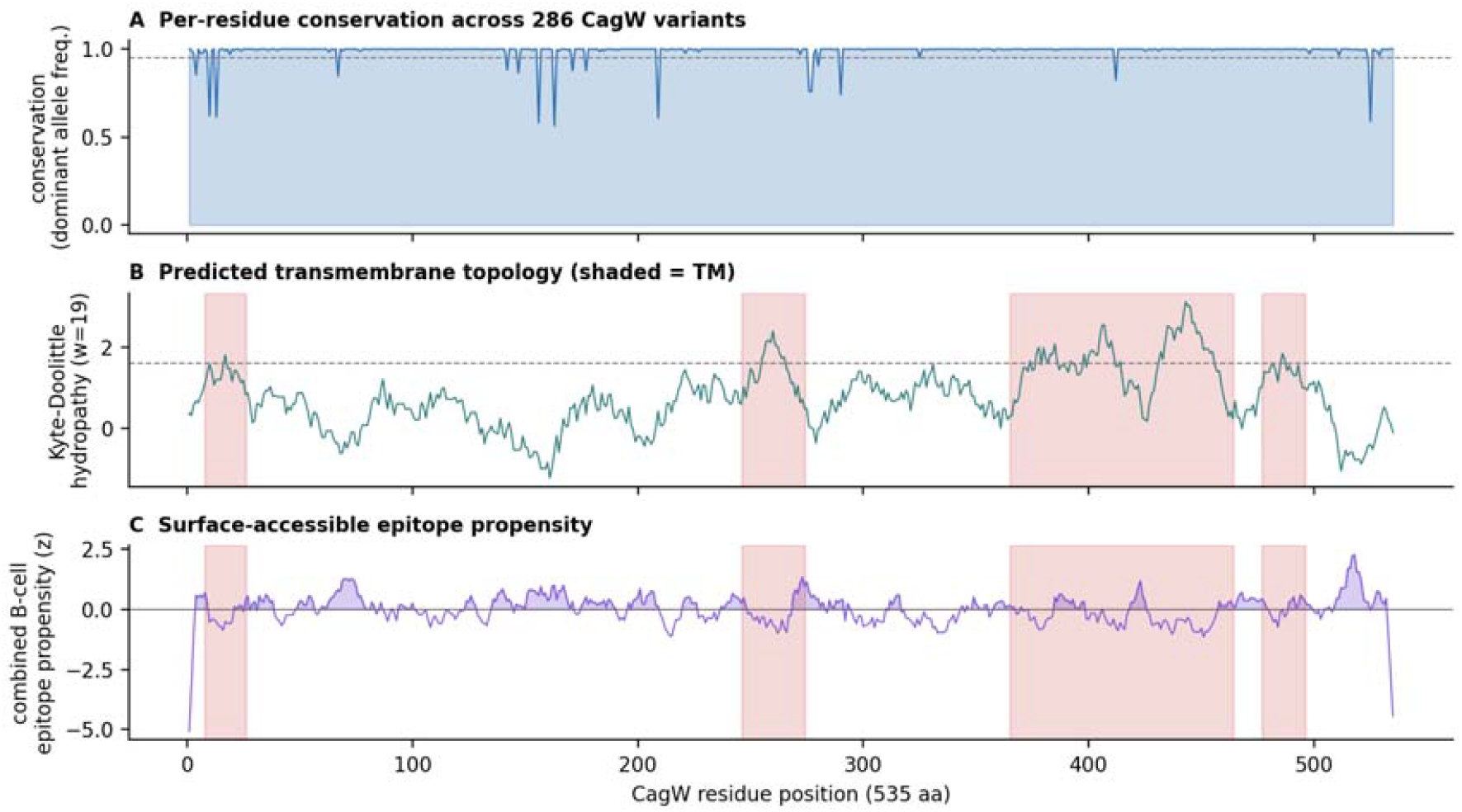
CagW antigen profiling across global *H. pylori* diversity (400 NCBI records; 286 unique variants). (A) Per-residue conservation (dominant-allele frequency). (B) Kyte-Doolittle hydropathy with predicted transmembrane segments shaded. (C) Combined linear B-cell epitope propensity score (mean z of four published scales) wit transmembrane segments shaded.

Combined linear B-cell epitope propensity, evaluated only in non-transmembrane windows, peaked sharply at the C-terminal cytosolic-soluble segment 505-530 (combined z up to 1.12; mean per-residue conservation in this window 0.996) and, secondarily, at the nucleotide-binding P-loop region 63-79 (z up to 0.88; conservation 0.988). The most conserved 15-mer windows of the entire protein lay at residues 194-208 (WLIGKEFTANKLQNN), 236-254 (VNLPDIIIGQGSYLDFV) and 301-319 (SFIAYAGFGYLYKIS), all with a dominant-allele frequency of exactly 1.0 (Figure 7C). Because these windows are simultaneously conserved and topologically accessible or proteasome-accessible, they constitute a rational epitope envelope for both antibody recognition and MHC-I presentation. For vaccine design, the four transmembran helices (31.4% of the sequence) imply that a secreted CagW-Fc fusion should focus on the soluble C-terminal (505-530) and P-loop (63-79) regions, or use isolated epitope peptides, rather than the full-length transmembrane protein.

### Integration: an immunological and metabolic niche that supports, and is shaped by, CagW vaccination

The four axes of evidence can now be read as a single system (Figure 1). *H. pylori* infection converts gastric epithelium into an NF-κB- and ATF4-dominated, Th17-recruiting, MHC-I-presentation-capable (with a non-significant trend toward MHC-I bias that did not survive correction) but MHC-II-restrained signalling platform, simultaneously depleting arginine and glutamine through polyamine synthesis and bacterial catabolism (Sections 3.2-3.4, 3.7). The mucosal cellular response to this niche is a stereotyped expansion of dendritic cells, macrophages and neutrophils with a modest T-cell and B-cell component (Section 3.5), within which a population of gastric CD8+ tissue-resident memory cells-shown for CagA-specific cells [21]-carries a locally acting cytotoxic program (Section 3.6); whether CagW-specific TRM are similarly induced remains to be established. A preventive mRNA-LNP vaccine is therefore working in an environment that is already rich in professional antigen-presenting cells and already skewed towards Th17-a situation that favours antigens delivered into the MHC-I pathway and adjuvanted by ionisable lipids, exactly as achieved by the CagW-Fc mRNA-LNP construct. Conversely, the metabolomic data predict that polyamine-driven arginine sequestration will restrain the nitric-oxide arm of the response; a vaccine that reduces bacterial burden should, on this logic, relieve this sequestration and thereby release a feed-forward antibacterial loop. (The epithelial signalling axis is inferred from human gastric cancer-cell-line bulk transcriptomes, whereas the cellular and metabolic axes derive from murine mucosa; this synthesis is therefore cross-species and requires validation in human tissue, e.g. GSE249874.) Finally, antigen-side constraints are satisfied: CagW is more than 98% identical across the modal length of global isolates, its most conserved regions are simultaneously topologically accessible or proteasome-accessible, and it is a constitutive, non-phase-variable component of the cagPAI that is present in the cagA+ strains that dominate gastric-cancer risk [6,7].

## Discussion

This study set out to replace intuition with evidence in the design of an mRNA vaccine against *H. pylori*. By mining rather than generating data, we were able to connect, in a single reproducible analysis, the transcriptional state of infected epithelium, the cellular composition of infected mucosa, the metabolic signature of infection and the population-genomic conservation of the vaccine antigen. Three findings are, to our knowledge, novel in this combination, and each has direct consequences for vaccine development.

First, the epithelial response is not simply ‘inflammatory’. It is a superposition of two separable programs-an NF-κB/chemokine module that is Th17-polarising and MHC-I-permissive, and an ATF4/integrated-stress module that is metabolic. The former provides the immunological substrate that a Th1/Th17 vaccine exploits; the latter predicts the metabolic cost of chronic infection and explains why infection depletes arginine and glutamine. The ATF4 signature (DDIT3, ATF3, CHAC1, TRIB3, PPP1R15A, ASNS, WARS1) has not, to our knowledge, been highlighted previously as a dominant feature of the *H. pylori* epithelial transcriptome, and it provides a mechanistic bridge to the metabolomics data in which epithelial ODC deletion accumulates arginine and reduces NF-κB-driven CXCL8 [16,18]. For vaccine development, this dual reading implies that an effective vaccine must both amplify the myeloid-recruiting chemokine program (which it will do, because mRNA-LNP self-adjuvancy engages the same NF-κB axis) and relieve the metabolic stress that restrains NO-mediated killing-an outcome that reduced bacterial burden is expected to deliver.

Second, single-cell re-analysis shows that the mucosal niche of chronic infection is not T-cell-poor, as is often assumed, but rather myeloid-rich and epithelially de-differentiated. Professional antigen-presenting cells more than double their share of captured cells, and B/plasma cells-the source of the mucosal sIgA that our CagW mRNA-LNP vaccine induces-increase almost twofold. At the same time the cells that maintain acid secretion and the mucus barrier (parietal, chief, pit and mucous-neck cells) are depleted by two-to five-fold. This combination is precisely the niche in which an intramuscular mRNA-LNP vaccine operating through a draining-lymph-node DC program plus mucosal homing can be expected to work, and it explains why gastric sIgA rather than serum IgG titres correlate best with protection in many mucosal infections (to be confirmed for this antigen *in vivo*). It also underscores that preventive vaccination should ideally occur before irreversible lineage loss.

Third, the CD8+ arm is not a secondary consideration. Gastric CD8+ CD69+ CD103+ cells induced by *H. pylori* express a genuine tissue-residency and cytotoxic program while losing all recirculation machinery; gastric CD8+ TRM cells are known to recognise CagA-derived peptide and control early infection [21], demonstrating that the cytotoxic-resident compartment is functionally engaged during *H. pylori* infection. Because mRNA delivered by LNP is presented efficiently on MHC-I, a CagW-based vaccine is mechanistically positioned to engage this compartment (pending human validation); CagW, unlike the secreted CagA, is membrane-associated and therefore continuously available for proteasomal processing, which may favour sustained MHC-I presentation.

Our antigen-side analysis addresses the central objection to monovalent *H. pylori* vaccines: strain diversity. By measuring conservation directly across 286 unique global CagW variants rather than relying on a single reference genome, we show that 96.4% of residues are ≥95% conserved and that the top epitope-bearing windows are 100% conserved. This places CagW in a different conservation class from hypervariable surface antigens such as BabA, SabA or OipA, and comparable to the most conserved core components of the T4SS. The four transmembrane helices predict limited antibody access to the membrane-spanning regions, but the C-terminal soluble segment (505-530) and the P-loop region (63-79) combine high conservation with high epitope propensity, and are therefore the segments that a structure-guided refinement of the antigen should preserve, pending HLA-binding and structural validation.

Several limitations must be acknowledged. The bulk transcriptomic contrast is in vitro and, although cell-line-blocked, does not capture the stromal and immune contribution of intact mucosa; the single-cell atlas is murine, and cellular composition was aggregated across 16,434 cells and tested at cell level, which over-states statistical power relative to the true n=8 sample-level design; sample-level effect sizes with confidence intervals should be reported as the primary statistic; the metabolomics datasets are targeted or partially annotated panels rather than deep untargeted profiles, and the MTBLS3235 arginine shift is an Odc-genotype main effect rather than a clean infected versus uninfected contrast; the CagW epitope analysis uses propensity scales rather than structure-based or HLA-binding predictions, so it nominates rather than validates epitopes, and the 96.4% residue conservation likewise includes fragmentary records and lacks a bootstrap confidence interval. Finally, the human gastric scRNA-seq resource GSE249874, which stratifies gastritis, intestinal metaplasia and gastric cancer by ^13^C-urea-breath-test-confirmed *H. pylori* status, was identified but not re-analysed here because its raw matrix exceeds 1.4 GB; it represents the natural next step for validating the mouse-derived composition findings in human tissue.

These limitations do not alter the central conclusion, but they define the experimental agenda: (i) confirm the epithelial ATF4 and polyamine signature in human biopsies stratified by infection status; (ii) test whether CagW mRNA-LNP immunisation relieves arginine sequestration in infected mucosa, thereby releasing NO-mediated killing; (iii) determine whether CagW-specific CD8+ tissue-resident memory cells are induced by the vaccine and whether, analogously to CagA-specific TRM [21], they control early colonisation; and (iv) evaluate CagW in combination with UreB or with a second cagPAI component to cover cagPAI-negative strains.

## Conclusions

By systematically re-analysing public next-generation sequencing, single-cell and metabolomics resources, we reconstruct the mucosal niche in which a vaccine against *H. pylori* must operate: an NF-κB/ATF4-driven, Th17-polarised, arginine- and polyamine-remodelled epithelium overlaid by an expanded dendritic-cell, macrophage and neutrophil compartment and by a locally fixed cytotoxic CD8+ tissue-resident memory population. Within this niche, CagW is a near-universally conserved, MHC-I-presentation-capable (with a non-significant trend toward MHC-I bias that did not survive multiple-testing correction), epitope-accessible antigen. These data provide a coherent systems-level rationale for prioritising CagW as the antigen of a CagW-Fc mRNA-LNP vaccine candidate, and they define the biomarkers (epithelial ATF4 activity, mucosal myeloid fraction, gastric sIgA, tissue-resident CD8+ phenotype and tissue arginine/polyamine ratio) by which next-generation formulations should be triaged.

## List of abbreviations

ATF4: activating transcription factor 4
ATF3: activating transcription factor 3
BH: Benjamini-Hochberg
cagPAI: cag pathogenicity island
CagW: VirB6-family component of the *H. pylori* type IV secretion system
Cbl: ^13^C-urea breath test
CPM: counts per million
CTH: cystathionine-γ-lyase
DC: dendritic cell
DEG: differentially expressed gene
FDR: false discovery rate
GEO: Gene Expression Omnibus
GRAVY: grand average of hydropathicity
HRI: heme-regulated inhibitor kinase
ISR: integrated stress response
LNP: lipid nanoparticle
LTB4: leukotriene B4
MHC: major histocompatibility complex
mRNA: messenger ribonucleic acid
NO: nitric oxide
ODC: ornithine decarboxylase
OR: odds ratio
pDC: plasmacytoid dendritic cell
PIGR: polymeric immunoglobulin receptor
scRNA-seq: single-cell RNA sequencing
sIgA: secretory immunoglobulin A
T4SS: type IV secretion system
TLR: Toll-like receptor
TRM: tissue-resident memory T cell
UPR: unfolded protein response.

## Author Contributions

Yunwang Zhao: investigation, formal analysis, writing-original draft. Hongxian Cheng: writing-review and editing, data curation. Shiyi Zhou: visualisation. Lei He and Chenlin Xiu: methodology, formal analysis. Baihai Huang: conceptualisation. Xiongxiong Lu: conceptualisation, methodology, writing-review and editing, supervision.

## Data Availability

All datasets re-analysed here are publicly available under the accessions listed in Table 1 (GSE295523, GSE224840, GSE216894, MTBLS3235, MTBLS2851, NCBI Protein). All analysis scripts, intermediate result tables and figures are provided as supplementary material.

## Ethics Approval and Consent to Participate

This study used only publicly deposited, de-identified data; no new human or animal work was performed.

## Acknowledgments

The authors thank the depositors of GSE295523, GSE224840, GSE216894, MTBLS3235 and MTBLS2851 for making their primary data publicly available.

## References

1. Polk DB, Peek RM. Helicobacter pylori: gastric cancer and beyond. Nat Rev Cancer. 2010;10:403–414.

2. Li Y, Choi H, Leung K, Jiang J, Wong MCS, Chan PKS, Leung WK. Global prevalence of Helicobacter pylori infection between 1980 and 2022: a systematic review and meta-analysis. Lancet Gastroenterol Hepatol. 2023;8:553–564.

3. Graham DY. Helicobacter pylori update: gastric cancer, reliable therapy, and possible benefits. Gastroenterology. 2015;148:719–731.

4. de Martel C, Georges D, Bray F, Ferlay J, Clifford GM. Global burden of cancer attributable to infections in 2018: a worldwide incidence analysis. Lancet Glob Health. 2020;8:e180–e190.

5. Zeng M, Mao XH, Li JX, Tong WD, Wang B, Zhang YJ, et al. Efficacy, safety, and immunogenicity of an oral recombinant Helicobacter pylori vaccine in children in China: a randomised, double-blind, placebo-controlled, phase 3 trial. Lancet. 2015;386:1457–1464.

6. Costa TRD, Felisberto-Rodrigues C, Meir A, Prevost MS, Redzej A, Trokter M, Waksman G. Secretion systems in Gram-negative bacteria: structural and mechanistic insights. Nat Rev Microbiol. 2015;13:343–359.

7. Backert S, Clyne M, Tegtmeyer N. Molecular mechanisms of gastric epithelial cell adhesion and injection of CagA by Helicobacter pylori. Cell Commun Signal. 2011;9:28.

8. Juhas M, Crook DW, Hood DW. Type IV secretion systems: tools of bacterial horizontal gene transfer and virulence. Cell Microbiol. 2008;10:2377–2386.

9. Chehelgerdi M, Doosti A. Effect of the cagW-based gene vaccine on the immunologic properties of BALB/c mouse: an efficient candidate for Helicobacter pylori DNA vaccine. J Nanobiotechnology. 2020;18:63.

10. Raudvere U, Kolberg L, Kuzmin I, Arak T, Adler P, Peterson H, Vilo J. g:Profiler: a web server for functional enrichment analysis and conversions of gene lists (2019 update). Nucleic Acids Research. 2019;47(W1):W191–W198.

11. Kyte J, Doolittle RF. A simple method for displaying the hydropathic character of a protein. J Mol Biol. 1982;157:105–132.

12. Parker JMR, Guo D, Hodges RS. New hydrophilicity scale derived from high-performance liquid chromatography peptide retention data: correlation of predicted surface residues with antigenicity and X-ray-derived accessible sites. Biochemistry. 1986;25:5425–5432.

13. Karplus PA, Schulz GE. Prediction of chain flexibility in proteins. Naturwissenschaften. 1985;72:212–213.

14. Emini EA, Hughes JV, Perlow DS, Boger J. Induction of hepatitis A virus-neutralizing antibody by a virus-specific synthetic peptide. J Virol. 1985;55:836–839.

15. Kolaskar AS, Tongaonkar PC. A semi-empirical method for prediction of antigenic determinants on protein antigens. FEBS Lett. 1990;276:172–174.

16. Chaturvedi R, Asim M, Hoge S, Lewis ND, Singh K, Barry DP, et al. Polyamines impair immunity to Helicobacter pylori by inhibiting L-arginine uptake required for nitric oxide production. Gastroenterology. 2010;139:1686–1698.

17. Gobert AP, Cheng Y, Akhtar M, Mersey BD, Blumberg DR, Cross RK, et al. Protective role of inducible nitric oxide synthase in Helicobacter pylori-induced gastritis in mice. Infect Immun. 2004;72:3673–3678.

18. Hardbower DM, Asim M, Luis PB, Singh K, Barry DP, Yang C, et al. Ornithine decarboxylase regulates M1 macrophage activation and mucosal inflammation via histone modifications. Proc Natl Acad Sci USA. 2017;114:E751–E760.

19. Pardi N, Hogan MJ, Porter FW, Weissman D. mRNA vaccines-a new era in vaccinology. Nat Rev Drug Discov. 2018;17:261–279.

20. Hou X, Zaks T, Langer R, Dong Y. Lipid nanoparticles for mRNA delivery. Nat Rev Mater. 2021;6:1078–1094.

21. Koch MRA, Gong R, Friedrich V, Alderlieste YAS, Vehreschild MJGT, Koch S, et al. CagA-specific gastric CD8+ tissue-resident T cells control Helicobacter pylori during the early infection phase. Gastroenterology. 2023;164:550–566.

22. Schenkel JM, Masopust D. Tissue-resident memory T cells. Immunity. 2014;41:886–897.

23. Son SY, Lee CH, Lee SY. Different metabolites of the gastric mucosa between patients with current Helicobacter pylori infection, past infection, and no infection history. Biomedicines. 2022;10:556.

24. Zang H, Wang J, Wang H, Guo J, Li Y, Zhao Y, et al. Metabolic alterations in patients with Helicobacter pylori-related gastritis: the H. pylori-gut microbiota-metabolism axis in progression of the chronic inflammation in the gastric mucosa. Helicobacter. 2023;28:e12984.

25. Keilberg D, Steele N, Fan S, Vagdama E, Southard GL, Mahe MM, et al. Gastric metabolomics detects Helicobacter pylori-correlated loss of numerous metabolites in both the corpus and antrum. Infect Immun. 2021;89:e00619–20.

26. Latour YL, Sierra JC, Finley JL, Asim M, Barry DP, Allaman MM, et al. Cystathionine gamma-lyase exacerbates Helicobacter pylori immunopathogenesis by promoting macrophage metabolic remodeling and activation. JCI Insight. 2022;7:e158336.

27. Barrett T, Wilhite SE, Ledoux P, Evangelista C, Kim IF, Tomashevsky M, et al. NCBI GEO: archive for functional genomics data sets - update. Nucleic Acids Res. 2013;41:D991–D995.

28. Haug K, Cochrane K, Nainala VC, Williams M, Li J, Chang J, et al. MetaboLights: a resource evolving in response to the needs of its scientific community. Nucleic Acids Res. 2020;48:D440–D444.

29. Kon E, Ad-El N, Hai I, Zarov I, Elia U, Cohen Y, et al. A single-dose F1-based mRNA-LNP vaccine provides protection against the lethal plague bacterium. Sci Adv. 2023;9:eadg1036.

30. Sayi A, Kohler E, Hitzler I, Genitsarioti I, Reusch C, Hardt WD, Muller A. The CD4+ T cell-mediated IFN-gamma response to Helicobacter pylori infection is essential for clearance and determines gastric cancer risk. J Immunol. 2009;182:7085–7101.

